# Disruption of IgA-mediated aggregation at weaning favors mucus encroachment by commensal bacteria

**DOI:** 10.1101/2025.07.25.665838

**Authors:** Kevin Simpson, Renaud Baillou, Tiphaine Le Roy, Axel Ranson, Marta Vazquez-Gomez, Delphine Sterlin, Guy Gorochov, Martin Beaumont, Karine Clément, Eric Clément

## Abstract

Disruption of the gut mucus barrier is a critical step in the development of infectious or chronic inflammatory diseases. However, there are no clear links between developmental stages, diet, and the mechanical and biochemical properties of mucus. The transition from suckling to weaning is a pivotal stage in the development of the mucus barrier in mammals, with significant implications for the health and morbidity of mammalian infants. Here, using a novel microfluidic device, we investigate the penetration and organizational properties of motile *Escherichia coli* bacteria at the mucus interface using purified intestinal mucus collected from cohorts of piglets before and after weaning. In weaned piglets, *E. coli* penetrate more than 100 ***µ***m into the mucus, a distance greater than the physiological thickness of the mucus layer *in vivo*. In contrast, for suckling piglets significant bacterial aggregation is observed at the interface, hindering the penetration process. Using the supernatant obtained from purified mucus of suckling piglets, we were able to restore bacterial aggregation in weaned piglet mucus and limit penetration. Interestingly, we also achieved the same result using purified human breast milk immunoglobulin A (IgA), which is known to promote bacterial aggregation. Our results emphasize the importance of mucosal immunoglobulin A (IgA) specificity in relation to the mother’s immunological history, which is primarily transmitted through breast milk and lost during weaning. This also might explain why the suckling/weaning transition is, among other issues, a critical window associated with a high incidence of gastrointestinal infections, before autologous IgA-mediated definite protection is acquired. Studying bacterial penetration in complex fluids using this new *in vitro* microfluidic device will pave the way for future research and the development of predictive tools for use in medical research trials.

## 1 Introduction

The intestinal mucus barrier is a critical component of mucosal immunity, acting as a dynamic and selective interface between the host epithelium and the luminal environment. It shields epithelial surfaces from microbial invasion, facilitates antigen sampling and immune education, while also allows nutrients absorption. This function is vital during early life, a period of immune and microbial co-development, where the structural and immunological integrity of the mucus layer shapes host–microbe interactions and immune homeostasis [1, 2]. The mucus layer consists primarily of the gel-forming mucin MUC2, secreted by goblet cells, along with immunoglobulin A (IgA), antimicrobial peptides, and other glycoproteins. These components assemble into a viscoelastic hydrogel that forms a physical and immunological barrier to microbes [3, 4]. From a physical perspective, the rheological properties of the mucus hamper bacteria penetration into the epithelium. It can be seen as a complex, high-viscosity fluid that exhibits yield stress, thus providing a solid, elastic resistance to penetration [5]. For a given mucins composition/structure, the rheological properties of mucus are influenced by (at least) the pH and the salt concentration encountered in the intestinal lumen, which can also be influenced by the luminal microbiome [6–9].

The suckling-to-weaning transition is a particularly vulnerable developmental period, characterized by a rapid dietary shift from maternal milk to solid feed, the loss of maternally derived immune factors such as secretory IgA, and increased exposure to microbial and dietary antigens. This abrupt transition can disrupt the coordinated maturation of the gut microbiota and mucosal barrier, thereby increasing susceptibility to infectious and inflammatory diseases later in life [10, 11]. Such disruptions have been shown to alter mucus composition and compromise barrier integrity [2, 12]. Importantly, this phase also represents a critical window during which dynamic interactions between diet, microbiota, and the developing immune system shape long-term metabolic and immune trajectories, a core principle of the Developmental Origins of Health and Disease (DOHaD) framework [13]. In agricultural settings, this period is also associated with a high incidence of gastrointestinal infections and impaired growth, representing a major concern in veterinary medicine [14]. For example, *Escherichia coli* infection induce post-weaning diarrhea, that can affect the production and lead to death [15]. More broadly, the translocation of *E. coli* through the mucosal barrier has been associated with the development of diseases [16]. Although motility in *E. coli* plays a critical role in overcoming the barrier [17], the properties underlying the swimming of bacteria in mucus are poorly understood. Beyond early development, altered mucus barrier function in the colon has been implicated in a range of chronic inflammatory conditions, including inflammatory bowel disease, type 2 diabetes, and obesity [18–20].

The mucus barrier is dynamic: it is continuously secreted by epithelial cells, degraded by the microbiota, and shed from the surface. Its coverage and properties can be modified in response to stimuli from intestinal contents, and dietary components play an important role in maintaining the mucus barrier [21–23]. In these contexts, increased microbial motility and barrier penetration are linked to epithelial stress and immune activation. Notably, Chassaing and colleagues demonstrated that individuals with metabolic disorders exhibit increased bacterial encroachment—defined as closer proximity of microbes to the intestinal epithelium—accompanied by a reduced mucus layer and increased inflammatory markers [24]. Furthermore, they showed that dietary emulsifiers, commonly found in processed foods, impair mucus structure and promote microbial translocation, contributing to the onset of colitis and features of metabolic syndrome in mice [24]. Moreover, western diets low in fiber but high in fat and sugar increase bacterial penetrability of the barrier [25], while high fat diet also reduces the thickness of the mucus layer [26].

In this study, we develop a novel and technically simple approach based on an *ex vivo* microfluidic system that enables real-time visualization and quantification of the penetration of flagellated bacteria in mucus samples under defined host and dietary conditions. We applied this platform to compare mucus from suckling and weaned piglets, a relevant model capturing dietary, microbial, and immunological transitions. This tool allows us to directly visualize and quantify across the suckling/weaning transition the significant differences in the penetration modes of motile *E. coli* into mucus. The model organism used in this work is a common commensal bacteria that is widely present in the gut microbiota, and is highly relevant to gut health [27]. We here emphasize the pivotal protective function of antigen-mediated bacterial aggregation during suckling and characterize its specificity to microbial penetration. We demonstrate how this protection can be rescued by transferring immunity from suckling to weaned piglets, or from maternal milk directly. This approach provides a tractable and quantitative tool to study mucus barrier function and its disruption across critical physiological windows.

## 2 Results

### 2.1 Enhanced mucus penetration by motile bacteria in weaned piglet

In order to study the effects of suckling/weaned transition and age-related changes on mucus and their impact in *E. coli* motility, we used samples of mucus extracted from the small intestine of suckling piglets, fed exclusively with milk for 21 days before sacrifice, and weaned piglets, which began to supplement their diet with solid feed at 21 days, completed weaning at 28 days and sacrificed at 35 days (Fig. 1a). These piglets were part of a recent study by Mussard et al. [28] that characterized the microbiota, metabolome, epithelial gene expression and mucosal morphology in these samples. Importantly, in order to remove large contaminants and small soluble molecules, the raw mucus samples were dialysed, washed, and resuspended in a buffer.

**Fig. 1.**
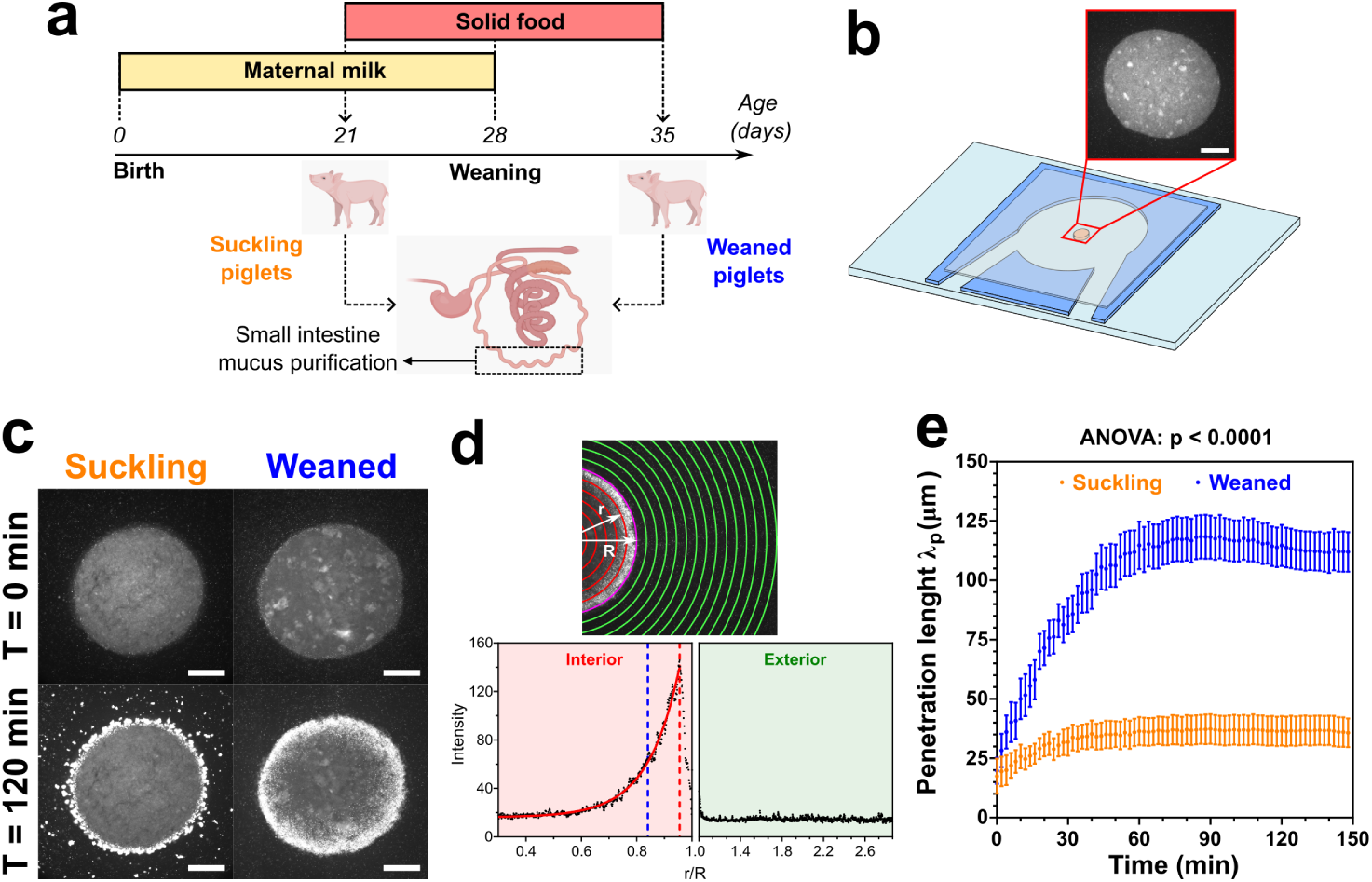
Suckling-to-weaning transition is associated with increased mucus permeability to motile bacteria. a) Mucus samples were collected from the small intestine of 21-day-old suckling piglets (orange) and 35-day-old weaned piglets (blue) (*n* = 7 per group). b) Schema of the *in-vitro* microfluidic system built to study the interface between a mucus droplet (orange and inset) and a *E. coli* suspension. Mucus droplets were placed in a chamber between a glass slide and a coverslip separated by a 100 *µ*m-thick spacer (blue). Scale bar 500 *µ*m. c) Initial (*T* = 0 min) and final (*T* = 120 min) images from a time-lapse experiment showing fluorescent *E. coli* (bright dots) in mucus droplets from suckling or weaned piglets. Scale bars 500 *µ*m. d) Quantification of bacterial penetration length *λp* in mucus droplets. The droplet boundary (magenta line) defines interior (red) and exterior (green) regions. Mean pixel intensity was calculated at each radial position *r* and plotted as a function of *r/R*, with *R* = droplet radius. *λp* (blue dashed line) was calculated by fitting an exponential decay function from the maximum value of the curve (red dashed line). e) Temporal evolution of *λp* in mucus from suckling (*n* = 7) and weaned (*n* = 7) piglets. Dots and error bars represent the mean of all samples per condition ± the standard error of the mean (SEM). Each sample was measured in quadruplicate. The result of a type III ANOVA performed on the linear mixed model derived from these data is shown on the graph, and the significant differences at each time were calculated by the estimated marginal means (EMMeans), corrected by Tukey’s method, and are shown in the supplementary table S1.

To monitor the dynamics of bacterial penetration in the piglet mucus samples, we designed an *ex-vivo* microfluidic assay to create an interface between a droplet of mucus and a bacterial suspension (Fig. 1b). We used the flagellated AD62 *E. coli* strain [29] (which constitutively expresses a green fluorescent protein) as a biological model for micro-swimmers performing run-and-tumble exploration kinematics. We placed 0.5 *µ*L mucus droplets inside our microfluidic system, which was then filled with the bacterial suspension. After sealing the device, the mean radius of the droplets inside the chambers was found to be 915.87 ± 96.39 *µ*m. Through time-lapse visualization of the mucus droplets, we observed that motile bacteria easily penetrate the mucus from weaned piglets, as observed by the bright dots (fluorescent bacteria) inside the mucus droplet at *T* = 120 min (Fig. 1c and S1). In contrast, mucus from suckling piglets rapidly halted bacterial movement, preventing penetration and restricting bacterial accumulation primarily to the periphery of the droplet (Fig. 1c and S1). All suckling mucus samples showed a similar behavior, except for the sample from piglet 4 (Fig. S1, Mucus 4). In this sample, bacteria easily penetrated inside mucus, and this sample had similar properties to the samples collected from weaned piglets.

To quantify the penetration of bacteria in the mucus samples in time, we used the fluorescence intensity radial density profiles of the time-lapse images. By fitting the intensity profiles in the interior mucus region by an exponential decay function starting at the maximum peak value (Fig. S2), we extracted the penetration length *λ_p_* of bacteria in mucus over time (Fig. 1d). We observed that the average value of *λ_p_* at *T* = 120 min in weaned mucus was greater than the average value of *λ_p_* for suckling mucus (*λ_p_* ≈ 115 *µ*m vs. *λ_p_* ≈ 37 *µ*m, p-value *<* 0.0001, by Mann-Whitney test, at *T* = 120 min) (Fig. 1e). Furthermore, significant differences between these two groups are observed as early as 18 min (Table S1). This experiment demonstrates that, in the case of suckling piglets, bacteria remain largely confined to the periphery of mucus droplets. In contrast, for weaned piglets, the bacteria penetrate deeply into the interior before becoming trapped. We also found that bacterial motility is essential for penetrating mucus. Motile *E. coli* were able to travel deep into the mucus of weaned piglets, whereas inert 1 *µ*m fluorescent particles could not infiltrate the layer (Fig. S3). Together, these findings reveal a significant shift in the biophysical properties of gut mucus during the transition from suckling to weaning, which affects the ability of motile bacteria to pass through this barrier.

### 2.2 Interface-driven bacterial aggregation in mucus from suckling piglets

To gain insights into the mechanisms underlying the breaching of the mucus barrier, we examined the spatial distribution of bacteria in the mucus droplets. In addition to deeper bacterial penetration in weaned piglet mucus, we observed the emergence of bacterial aggregates at the boundary of mucus droplets from the suckling piglets (Fig. 1c at *T* = 120 min). In the case of suckling piglet mucus, the bacterial aggregates formed at the boundary of the droplet and also extended outward into the surrounding medium (Fig. 2a and S1). By contrast, bacterial aggregation did not occur outside the mucus droplets from weaned piglets, and aggregates were essentially found inside them (Fig. 2a and S1). To quantify these observations, we measured the fraction of the area occupied by bacterial aggregates larger than 27 *µ*m^2^ at *T* = 120 min minutes in different regions of the mucus droplets (Fig. 2b). This threshold was chosen as a minimum size of around 10 bacteria, which allows for significant determination above noise for a cluster. For suckling piglets, the proportion of the area occupied by the aggregates peaked in the region immediately outside the droplet boundary, and decreased abruptly to almost zero at 80% of the radius inside the droplet. On the other hand, the maximum concentration of aggregates in mucus from weaned piglets was found in the first region inside the droplets. In this samples, aggregates could even be found as deep inside as 40% of the droplet radius. These observations suggest that mucus from suckling and weaned piglets may exhibit functionally divergent properties.

**Fig. 2.**
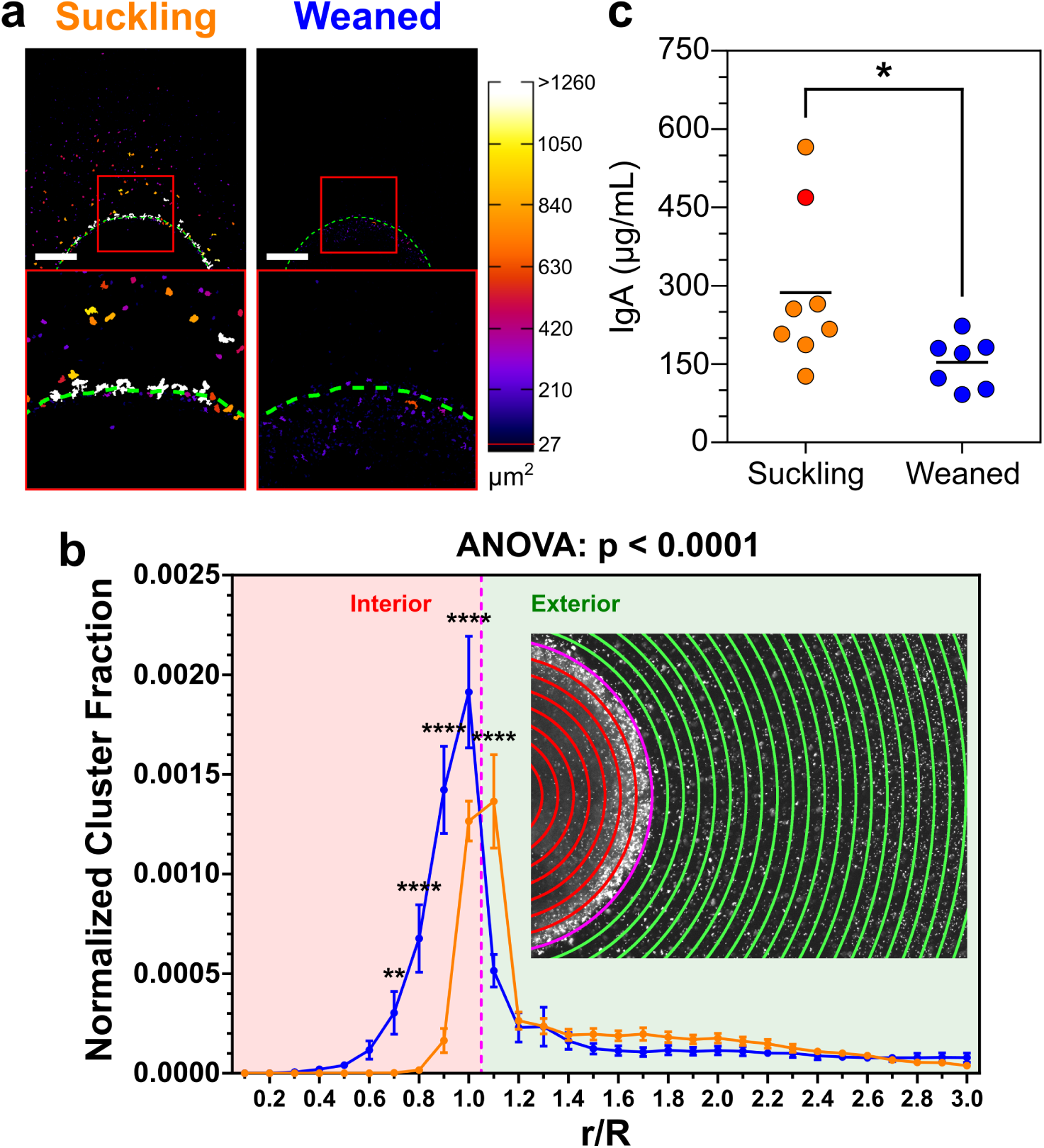
Mucus from suckling piglets promotes bacterial aggregation at the mucus/bac-terial suspension interface. a) Examples of binarized images of mucus droplets from suckling and weaned piglets at *T* = 120 min. Bacterial aggregates are color-coded according to their area. Green dashed lines delineate the mucus edges. Bottom panels show enlarged views of the red boxes in top panels. Scale bars 500 *µ*m. b) Quantification of the normalized fraction of aggregated bacteria at *T* = 120 min across defined sub-regions of mucus droplets from suckling (orange, *n* = 7) and weaned (blue, *n* = 7) piglets. Data represent the mean ± SEM of all piglets per group, measured in quadruplicate. Inset: Mucus droplet with overlaid radial zones used for quantification of the fraction of aggregated bacteria inside (red lines) and out side (green lines) the droplet. Magenta lines: Middle mucus boundary. The result of a type III ANOVA per-formed on the linear mixed model derived from these data is shown on the graph. Significant differences at each *r/R* were calculated by the estimated marginal means (EMMeans), and corrected by Tukey’s method (∗∗: p ≤ 0.01, ∗ ∗ ∗∗: p ≤ 0.0001, Table S2). c) Immunoglobulin A (IgA) concentration in purified mucus collected from suckling and weaned piglets. Each point represents the IgA concentration of an individual sample, and horizontal black lines indicate group means. Red point highlights the IgA concentration of the suckling piglet 4. Result from non-parametric, unpaired two-tailed Mann-Whitney test is show on the graph (∗: p *<* 0.05, *α* = 5%).

The presence of immunoglobulin A (IgA) in breast milk is thought to confer protective effects against gastrointestinal diseases [30]. In mucus, IgA is involved in agglutination, entrapment, and clearance of microbes [31], but its concentration decreases during the weaning period [32, 33]. Thus, the reduced IgA levels in weaned mucus may impair its ability to aggregate motile bacteria, thereby contributing to the increased bacterial penetration observed. Quantification of IgA levels in the purified mucus samples revealed, on average, a higher concentration in mucus from suckling piglets than in mucus from weaned piglets (Fig. 2c). Although the difference was statistically significant (p = 0.0140), it was largely driven by two samples, thereby weakening the overall correlation between IgA concentration and the presence of bacterial aggregates. Notably, one of the samples with the highest IgA levels—mucus 4 (Fig. 2c, red point)—did not promote bacterial aggregation and exhibited a penetration length (*λ_p_*) comparable to that of weaned piglets (Fig. S1). These findings suggest that IgA concentration alone is insufficient to account for the variability in aggregation behavior across samples. We therefore propose that IgA specificity, rather than total abundance, may be a critical factor in mediating bacterial aggregation at the mucus boundary in suckling piglets.

### 2.3 The supernatant from suckling piglet mucus and breast milk IgA restore barrier function in weaned piglet mucus

To determine whether components present in the mucus from suckling piglets con-fer resistance to bacterial penetration, we separated mucus samples from a suckling and a weaned piglet into pellet and supernatant fractions (Fig. 3a). We first tested whether these soluble fractions affected bacterial aggregation by supplementing bac-terial suspensions with the supernatants of the samples. As controls, we supplemented the bacterial suspensions with motility buffer or with purified IgA from human breast milk (30.4 *µ*g/mL). After 120 minutes of incubation, we observed increased bacterial aggregation only in suspensions supplemented with suckling mucus supernatant or with purified IgA (Fig. 3b). These results suggest that elements present in the super-natant of suckling mucus promote bacterial clustering, similar to the effect of purified IgA from breast milk. These elements appear to be absent (or present at insufficient levels) in the supernatant of weaned mucus.

**Fig. 3.**
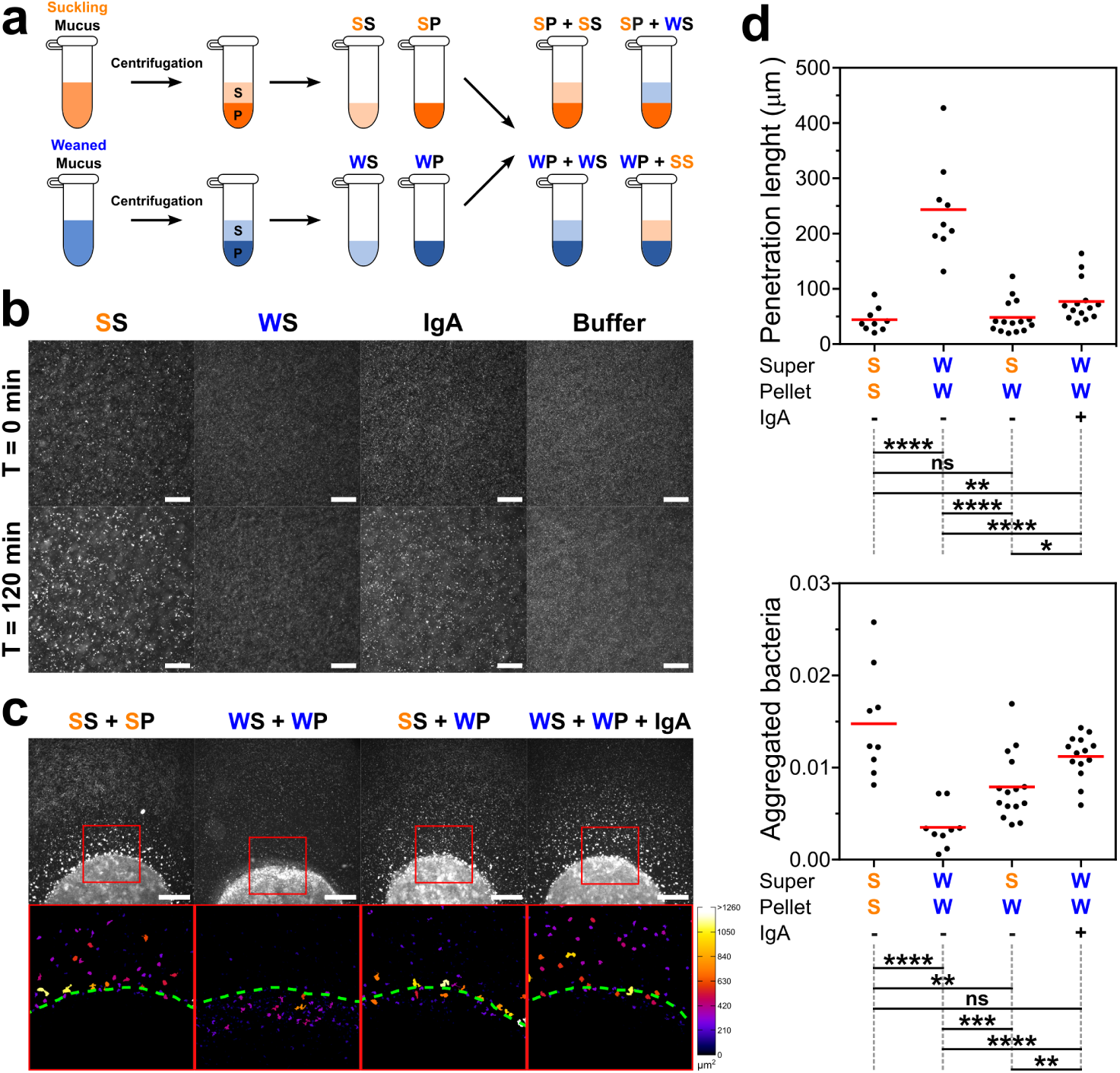
The supernatant from suckling mucus and IgA promotes bacterial aggregation and limits bacteria penetration into mucus. a) Protocol used to separate the supernatant and pellet of purified mucus from a suckling and a weaned piglet (*n* = 1 per condition), and to generate samples with different combinations of pellet and super-natant. For panels a–c, abbreviations are as follows: S: supernatant, P: pellet, SS: Suckling Supernatant, SP: Suckling Pellet, WS: Weaned Supernatant, WP: Weaned Pellet. b) Bacterial aggregation after the addition of mucus supernatant from suckling or weaned piglets, puri-fied IgA (30.4 *µ*g/mL), or motility buffer to a bacterial suspension. Images were acquired at *T* = 0 min (top) and *T* = 120 min (bottom). Scale bars 100 *µ*m. c) Representative images of droplets from reconstituted mucus obtained by mixing supernatant and pel-let from suckling and weaned mucus. Droplets were introduced into the microfluidic system, exposed to a bacterial suspension, and imaged after 120 min. Bottom panels show enlarged, binarized views of the red boxes in the top panels, with bacterial aggregates color-coded according to their area. Green dashed lines indicate mucus boundaries. Scale bars 500 *µ*m. d) Quantification of penetration length (top) and total fraction of aggregated bacteria outside mucus (bottom) in mucus droplets composed of different supernatant/pellet combinations from Suckling (S) or Weaned (W) mucus, with (+) or without (-) IgA supplementation. Each dot represents and individual experiment. Red horizontal lines denote group means. The table below each graph show the statistical differences based on Non-parametric, unpaired two-tailed Mann-Whitney test (*α* = 5%). ns (not signifi-cant): p *>* 0.05, ∗: p ≤ 0.05, ∗∗: p ≤ 0.01, ∗ ∗ ∗: p ≤ 0.001, ∗ ∗ ∗∗: p ≤ 0.0001.

We next reconstituted new mucus samples by combining pellet and supernatant fractions from suckling and weaned mucus in different configurations (Fig. 3a). After *T* = 120 min of incubating droplets from these samples with a bacterial suspension in our microfluidic setup, we observed that droplets containing suckling supernatant or added IgA produced bacterial aggregates outside the droplets, and showed a reduced accumulation of bacteria inside them (Fig. 3c). The quantitative analysis of these observations (Fig. 3d) confirmed that: *i*) the fully suckling (suckling supernatant + suckling pellet, SS-) and weaned (weaned supernatant + weaned pellet, WW-) combinations behave as their native condition, with low penetration and high aggregation for fully suckling and high penetration and low aggregation for fully weaned; *ii*) the addition of IgA to a weaned mucus (weaned supernatant + weaned pellet + IgA, WW+) partly restored its barrier properties, decreasing the penetration of bacteria and promoting their aggregation; and *iii*) replacing the supernatant of weaned mucus with that from suckling mucus (suckling supernatant + weaned pellet, SW-) also partly restored its barrier function, with a similar effect to IgA supplementation. On the other hand, replacing the suckling supernatant in a suckling mucus with weaned supernatant did not impair the barrier function of the suckling mucus (weaned supernatant + suckling pellet, WS-in Fig. S4). The low penetration and a high aggregation observed in this condition suggest that the elements present in the supernatant are also retained in the pellet, probably embedded within the mucin network. For penetration length, the following comparisons were significant: WW-vs. SS-(p *<* 0.0001), WW+ vs. SS-(p = 0.0085), SW-vs. WW-(p *<* 0.0001), WW+ vs. WW-(p *<* 0.0001), and WW+ vs. SW-(p = 0.0105), while SW-vs. SS-was not significant (p = 0.9742). For the total fraction of aggregated bacteria outside the droplets significant differences were observed for WW-vs. SS-(p *<* 0.0001), SW-vs. SS-(p = 0.0017), SW-vs. WW-(p = 0.0007), WW+ vs. WW-(p *<* 0.0001), and WW+ vs. SW-(p = 0.0068), while WW+ vs. SS-was not significant (p = 0.2003).

The spatial distribution of bacteria within these mucus droplets revealed that the addition of suckling supernatant or purified IgA to weaned mucus increased the fraction of area occupied by bacterial aggregates just outside the droplet boundary, while simultaneously causing a marked reduction in aggregates within the interior of the droplets (Fig. S5). Altogether, these findings suggest that soluble factors present in suckling mucus are sufficient to restore the ability of the mucus barrier to aggregate and immobilize motile bacteria, with secretory IgA, likely derived from maternal milk, emerging as a strong candidate underlying this phenotype.

## 3 Discussion

The transition from a milk-based to a solid diet is recognized as a critical stressor that affects intestinal physiology [34, 35], leading to impaired barrier function and increased susceptibility to infection [14, 36, 37]. Alterations in mucosal barrier properties and immune exclusion mechanisms during this sensitive period may have lasting effects on host-microbiota interactions and disease susceptibility later in life; thus, this weaning transition can be considered through the lens of the developmental origins of health and disease (e.g., DOHaD) framework [38].

In our study, we used mucus extracted from pigs. This animal model has emerged as an excellent means of studying human gut physiology due to the similarities between the two species in terms of nutrition, intestinal structure, functionality and microbiota [39–41]. Using a newly developed microfluidic platform, our study reveals significant alterations in mucus barrier properties across the suckling-to-weaning transition. These alterations have important implications for gut protection in early life and we reveal here an explicit mechanistic pathway susceptible to shed new lights on this critical period with possible therapeutic assays directions. Examining interaction between motile bacteria and mucus droplets we show that mucus from suckling piglets significantly restricts bacterial penetration and promotes bacterial aggregation at the droplet boundary. In contrast, mucus from weaned piglets allows deeper bacterial infiltration, accompanied by a loss of aggregation behavior. Importantly, we observed that *E. coli* penetrated more than 100 *µ*m into the mucus extracted from weaned piglets—a distance that exceeds the physiological thickness of the intestinal mucus layer in pigs [42, 43]. These findings point to a functional shift in the biophysical and immunological properties of mucus after weaning, that could contribute to digestive disorders observed in piglets at this stage.

A central question emerging from these results concerns the mechanisms underpinning this shift. Among the most plausible candidates, immunoglobulin A (IgA), particularly maternal secretory IgA, is a well-established effector of mucosal immunity and is frequently implicated in regulating microbial positioning at mucosal surfaces [30, 44, 45]. Through antigen-specific binding, IgA promotes bacterial aggregation and immune exclusion (e.g., the agglutination, mucus entrapment, and removal of pathogens), thereby restricting microbial diffusion and reducing contact with the epithelium [46]. Here, we observed a mean reduction in total IgA concentrations in weaned mucus compared to suckling mucus, which is expected as intestinal IgA is mainly provided by maternal milk in early life [33]. However, the variability in total IgA levels across samples, coupled with the absence of a consistent link with bacterial aggregation, prevent drawing definitive conclusions from these measurements alone. Notably, a suckling mucus sample with high IgA concentration failed to promote bacterial aggregation, further suggesting that IgA specificity, rather than concentration alone, plays a critical role in its barrier function. Although glycosylation in IgA and its secretory components should not be overlooked as it can promote bacterial aggregation independently of IgA specificity [47, 48], our observations aligns with recent studies showing that polyreactive or low-affinity IgA, which predominantly arise from T cell–independent responses [49, 50], are less effective at immobilizing bacteria within mucus. In contrast, high-avidity, antigen specific IgA promote robust bacterial aggregation through cross-linking, thereby enhancing immune exclusion at mucosal surfaces [51–54]. These findings suggest that while secretory IgA is abundant in mucus, its protective efficacy to agglutinate and clear pathogens relies on its ability to specifically recognize and bind to bacterial antigens.

Specific IgA acts as a molecular cross-linker to form multi-cell complexes that stick in mucus, whereas nonspecific or low-affinity IgA (even if abundant) cannot generate these aggregates on its own [55]. Interestingly, secretory IgA affinity in mucus increases over time with antigen exposure [56–58]. In suckling piglets, IgA is primarily derived from maternal milk, which contains highly matured, “antigen-experienced” antibodies [59]. In contrast, IgA present in weaned mucus is produced endogenously by the developing immune system of the piglet [60], and likely represents a population of antibodies with lower affinity due to limited antigen exposure and immune maturation. This difference in IgA affinity could underlie the reduced ability of weaned mucus to agglutinate our motile *E. coli* and limit their penetration. This hypothesis is further supported by our supernatant exchange experiments. Indeed, supplementation of weaned mucus with either suckling mucus supernatant or purified human breast milk IgA restored bacterial aggregation and significantly reduced penetration. These findings indicate that soluble immune components, likely including secretory IgA with appropriate antigen specificity, are critical for the functional integrity of the mucus barrier. However, it is important to note that other maternal or mucus-derived soluble factors not tested in this work, such as secretory IgM, antimicrobial peptides, glycosylated oligosaccharides, or low-molecular-weight mucin fragments [61, 62], may also contribute to the enhanced protective properties observed in suckling mucus independently of the bulk mucin network.

The observed difference in bacterial motility between mucus types adds another layer of interpretation. While motile *E. coli* could readily traverse weaned mucus, passive particles of similar size were excluded, underscoring the active role of motility in barrier breach. The empirical parameter *λ_p_*, quantifying penetration depth, may serve as a useful proxy for evaluating mucus barrier “quality” in future studies. Importantly, these results are consistent with prior work showing that early weaning impairs mucosal development and reduces mucus protective capacity [36]. From a physiological standpoint, breast milk appears to support both structural (mucin network) and immune (IgA-mediated) elements of the barrier [63]. Disruption of this support at weaning likely contributes to increase pathogen exposure and inflammation. This highlights the suckling-to-weaning transition as a particularly vulnerable window during which the rapid loss of maternal immune factors and structural changes in mucus can compromise gut barrier integrity and increase susceptibility to infection—an issue frequently observed in farming conditions [64], where early-life gastrointestinal disturbances can impair animal growth and represent a major concern in veterinary medicine [65]. Our findings have thus broader implications beyond swine physiology.

The microfluidic model developed here offers a powerful and tractable system to investigate bacteria–mucus interactions under controlled conditions, allowing dissection of the relative contributions of physical, immunological, and microbial factors. It can be adapted to other animal models, human samples, dietary conditions, or bacterial strains and including commensals and pathogens, hereby opening new avenues for studying host–microbiota interfaces. From a translational perspective, this work supports the rationale for strategies aimed at reinforcing the mucus barrier during vulnerable periods such as weaning. This could include nutritional interventions [66], microbiota-targeted approaches [26], or engineered IgA formulations [67] to restore aggregation and exclusion functions in immature or compromised guts.

## 4 Methods

### 4.1 Mucus sampling

#### 4.1.1 Mucus collection and purification

The piglets used in this study originated from the cohort described in [28]. The extraction of raw intestinal mucus was performed following the protocol described in [68]. The purification procedure was adapted from [69] using the equipment available in our laboratory.

#### 4.1.2 Extraction of raw mucus

As it was not always possible to obtain the small and large parts of the intestine, in most cases only the small intestine (from the duodenum to the ileum) was used to extract the raw mucus. However, there are a few samples that include mucus from the available large intestine; mainly from the descending colon and the centrifugal turns of the ascending colon. Immediately after the piglets were sacrificed, the luminal content of the fresh intestines was gently flushed with pure water. The intestines were then longitudinally opened, washed twice with water, and blotted dry. Mucus was collected by carefully scraping the inner surface with a sterile cell scraper (Falcon^®^Cell Scrapers #353089) to avoid scraping the tissue. The raw mucus extracted in this way was frozen before shipment.

#### 4.1.3 Purification of mucus

The raw mucus was first diluted 1/5 in sterile water and filtered through a standard sterile compress to remove large contaminants such as tissue debris. To eliminate small soluble molecules, including nutrients, the filtrate was dialyzed for 3 days at 4 *^◦^*C in deionized water using membranes with a 12–14 kDa molecular weight cut-off (the size of secretory IgA is around 300-400 kDa [70]), with daily water bath change. Further purification was achieved via centrifugal ultra-filtration (pore size: 50 kDa; 20,000 g at 4 *^◦^*C for 30 min). The retained material was collected by scraping the filter. This step was repeated on the supernatant until no further mucus could be recovered. Mucus was stored in 2 mL Eppendorf tubes at −20 *^◦^*C. Across all piglets, the final purified mucus had a stable pH (6.35 ± 0.10) and dry mass content (8.66 ± 0.77%).

#### 4.1.4 Ethics approval

All animal experimentation procedures were approved by the local ethics committee (N°TOXCOM/0136/PP) in accordance with the European directive on the protection of animals used for scientific purposes (2010/63/EU) [28].

### 4.2 Bacterial strains

All the experiments were performed using the *E. coli* AD62 [29], which is derived from the K-12 strain AB1157. This strain contains the plasmid pWR21, which confers resistance to ampicillin and also allows constitutive expression of a green fluorescent protein (eGFP) for the fluorescent imaging of the bacterial body.

### 4.3 Mucus penetration assay

#### 4.3.1 Preparation of the bacterial suspension

Bacteria pre-cultures were grown overnight in a shaking incubator at 30 *^◦^*C and 200 rpm/min in 4 mL of liquid Lysogeny broth (LB) (VWR Life Science) supplemented with 100 *µ*g/mL ampicillin. The overnight cultures were diluted 1:100 in Tryptone Broth (TB, 10 g/L Tryptone, 5 g/L NaCl) supplemented with 100 *µ*g/mL ampicillin and incubated at 30 *^◦^*C until optical density at 600 nm (OD_600_) 0.3 −0.5. The bacteria were centrifuged at room temperature at 3290 g per 5 minutes and the pellet was carefully re-suspended in 1 mL of Berg’s Motility Buffer X2 (BMB X2, 20 mM Phosphate Buffer (PB 100 mM: 61.5 mM K_2_HPO_4_ + 38.5 mM KH_2_PO_4_), 7.8 g/L NaCl, and 0.2 mM EDTA) supplemented with 25 g/L L-Serine (Sigma). The OD_600_ of the bacterial suspension was adjusted to 0.2, and then diluted 1:1 in Percoll (MP Biomedicals) solution to avoid sedimentation and achieve a non-buoyant bacterial suspension. When needed, 1 *µ*m fluorescent beads (Fluoro-max, Thermo) were added to test for the penetration of passive objects in mucus.

#### 4.3.2 Interface formation and Time-Lapse

To create an interface between mucus samples and bacteria, we designed a microfluidic device consisting of a “pool” of 14 mm in diameter and 100 *µ*m deep. To create this device, we cut a 14 mm diameter circle in a 100 *µ*m-thick double-sided adhesive tape (Tackotec) and stick it to a glass slide. Using a scalpel, we cut the tape to create the inlet and outlet. Before sampling, tubes with purified mucus were thawed for 1 h at room temperature in a rotator, and mucus solutions were homogenized by vortexing. A 0.5 *µ*L mucus droplet was carefully placed in the center of the pool, which was immediately covered with a glass coverslip. The pool was then filled by capillarity with the bacterial suspension in BMBx2:Percoll 1:1, slowly letting it come in contact with the mucus droplet. The inlet and outlet of the pools were then sealed with nail polish (Minute Quick-Finish Dry Varnish, Mavala) to prevent evaporation. The devices were placed immediately under an inverted epifluorescence microscope (Zeiss Observer Z1) equipped with a 5X objective to automatically record a time-lapse at room temperature using a motorized stage. Images (2048×2048 pixels) were captured every 2 minutes using a Hamamatsu ORCA Fusion camera and Colibri blue LED 30% (Zeiss Colibri 7). To capture the dynamics of the bacterial penetration, emergence of bacterial aggregates, and the behavior of the bacterial population on the same image, time-lapse images were acquired by focusing on a region covering approximately one-third of each mucus droplet. This allowed simultaneous observation of bacterial behavior both inside and outside the mucus droplet. When needed, a two-color LED light source and a dichroic image splitter (Hamamatsu) were used to simultaneously record the fluorescence of bacteria (green) and beads (red).

### 4.4 Mucus Supernatant

#### 4.4.1 Extraction of mucus supernatant

The extraction of the supernatant from purified mucus was performed on one sample from each group: suckling piglet 10 and weaned piglet 15. The purified mucus were centrifuged at room temperature at 970 g per 10 min, and the supernatants were transferred to Eppendorf tubes and stored at −20 *^◦^*C until their utilization. To test the ability of these supernatants to promote the aggregation of bacteria, 90 *µ*L of bacterial suspension in BMBx2:Percoll 1:1 were mixed with 10 *µ*L of the supernatants. As controls, we supplemented 49 *µ*L of the bacterial suspension with 1 *µ*L of 1.52 mg/mL immunoglobulin A (IgA) purified from human breast milk (positive control for aggregation of bacteria) and BMBx2:Percoll 1:1 (negative control for the aggregation). The purified IgA was provided by D. Sterlin and G. Gorochov laboratory. Fresh breast milk of healthy lactating women was obtained from the Lactarium regional d’Ile de France (Hopital Necker Enfants-Malades) and approved by the ethical committee (Aves CENEM 2020-V4 Biocol_lactothèque_V1.0_20190408). Secretory IgA was purified by using peptide M (InvivoGen) as described in [71]. The samples were then loaded into the microfluidic device consisting of a pool (in absence of mucus droplets), and images were acquired at *T* = 0 min and *T* = 120 min.

#### 4.4.2 Weaned and Suckling mucus supernatant exchange

To exchange the supernatant between mucus samples, 50 *µ*L of mucus from weaned and suckling piglets were centrifuged at room temperature at 970 rcf per 10 min. 15 *µ*L of supernatants were removed from each tube and transferred to Eppendorf tubes. The remaining pellets were supplemented with 15 *µ*L of the appropriate supernatant, in such a way to create the different combinations of pellet-supernatant (weaned pellet + weaned supernatant, weaned pellet + suckling supernatant, suckling pellet + suckling supernatant, and suckling pellet + weaned supernatant). The tubes were vortexed to allow the mix of pellet and supernatant. For the supplementation of mucus with IgA, 20 *µ*L of a mucus sample of weaned pellet + weaned supernatant was centrifuged at room temperature at 970 rcf per 10 min, 2 *µ*L of supernatant were replaced with 2 *µ*L of 1.52 mg/mL IgA, and finally vortexed. 0.5 *µ*L droplets of these samples were placed in the center of the pool as before, which was then filled with the bacterial suspension in BMBx2:Percoll 1:1, and images of these droplets were taken after two hours of incubation at room temperature.

### 4.5 Immunoglobulin A quantification in purified mucus

Purified mucus samples were thawed on ice and diluted 1:2000 in Tris-buffered saline (TBS) supplemented with 1% Bovine serum albumin (BSA) and 0.5% Tween 20. Porcine IgA were quantified by ELISA using Goat anti-Pig IgA Antibody (diluted 1:100, cat# A100-102A, Bethyl) for capture and Goat anti-Pig IgA Heavy Chain Antibody HRP Conjugated (diluted 1:50 000, cat# 100-102P, Bethyl) for detection. Revelation was performed with the BD OptEIA™ TMB Substrate Reagent Set (cat# 555214, BD Biosciences). A standard curve for absolute quantification was obtained with Pig IgA (swine non-immune, isotype control) (cat# 0015-20017-4-1-25, Gentaur).

### 4.6 Image analysis

The processing and analysis of the images was performed using the Fiji distribution of ImageJ [72]. Before the analysis, the background was removed on each image using the Subtract Background command.

#### 4.6.1 Penetration length

For the computation of the penetration length, the droplet mucus boundaries were manually defined using the Multi-Point Tool. This boundary divides the images into mucus exterior and mucus interior. For time-lapse experiments, the mucus boundaries were defined using the images at *T* = 0 min. Next, the radial intensity profiles were calculated for each image. An initial circle was manually drawn using the Oval Tool to approximate the position and overall shape of the droplets. Using this initial circle, the images were divided into a series of concentric rings centered on the mucus droplet an spaced 1 pixel apart, enabling fine-resolution sampling of the radial intensity. At each radial position *r*, the intensity was computed as the sum of pixel values along the corresponding circle, and then divided by the number of pixels within the image bounds at that radius. The resulting normalized intensities were plotted as a function of normalized radial distance *r/R*, where *R* is the droplet radius. To quantify the penetration length, an exponentially decaying function *y* = *Ae^−λx^* + *B* was fitted to the portion of the curve of the mucus interior extending inward from the maximum value of the radial intensity profile. Thus, 1*/λ* defines the penetration length *λ_p_*, which is the characteristic length that describes the decay of the bacterial concentration after its maximal.

#### 4.6.2 Aggregation of bacteria

To quantify the fraction of aggregated bacteria, images were first binarized using the Triangle method via the Make Binary Tool. Based on the droplet mucus boundaries and concentric rings computed for the calculation of the penetration length, we defined two circular boundaries around the mucus droplets: the Inner mucus boundary, corresponding to the largest circle that fits inside the mucus droplet boundary (i.e., the maximal inscribed circle that lies entirely within the droplet boundary), and the Outer mucus boundary, which corresponds to the smallest circle that fully encloses the mucus droplet boundary (i.e., a circumscribed circle that fully contains the mucus droplet boundary). The middle mucus boundary corresponds to the concentric rings in the middle of these two boundaries, and it divides the images in two regions: the interior and the exterior region of the droplets. Each region was further divided into concentric annular subregions. To normalize measurements of droplets of varying sizes across experiments, the interior region was divided into 10 concentric subregions of equal radial step size. The exterior region was divided using the same radial step size as in the interior. In each subregion, the fraction of area occupied by bacterial aggregates was calculated by measuring the area of white pixels (aggregated bacteria) using the Analyze Particles Tool, and then dividing by the total area of that region. Only aggregates larger than 27 *µ*m^2^m, which corresponds to the size of an aggregate containing approximately 10 bacteria, were considered for this analysis. To account for differences in the area of the subregions between experiments, the fraction of area occupied by bacterial aggregates was normalized by the the radial step size. Finally, we plotted the normalized cluster fraction 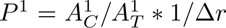 as a function of the radial step. The total fraction of aggregated bacteria was calculated by dividing the total area of white pixels in the Exterior region by the total area of that region. To illustrate the difference in size and position of bacterial aggregates in the binarized images, we used the ROI Color Coder plugin.

### 4.7 Statistics and bioinformatic analyses

Mucus was initially extracted from 8 suckling (samples 1, 3, 4, 9, 10, 17, 18, 20) and 7 weaned piglets (samples 5, 6, 8, 14, 15, 23, 24). The quantification of IgA was performed on all samples (*n* = 8 for suckling and *n* = 7 for weaned). However, suckling mucus 17 was excluded from further experiments due to failure to form stable droplets in the microfluidic setup. Therefore, the final dataset used for quantifying bacterial penetration and aggregation consisted of *n* = 7 suckling and *n* = 7 weaned mucus samples. For the weaned and suckling supernatant exchange experiments, we selected the mucus from suckling piglet 10 and weaned piglet 15. The number of replicates for each supernatant/pellet combination ranged between 9 and 15: *n* = 9 for suckling supernatant + suckling pellet (SS-), *n* = 9 for weaned supernatant + weaned pellet (WW-), *n* = 15 for suckling supernatant + weaned pellet (SW-), *n* = 11 for weaned supernatant + suckling pellet (WS-), and *n* = 14 for weaned supernatant + weaned pellet + IgA (WW+).

Statistical tests used are indicated in the legend to each figure. Statistical analyzes for curve differences were performed with R (version 4.3.3) (https://www.R-project.org/). A linear mixed model was generated to assess the effect of group, time or r/R and their interaction on the measured variable, followed by a type III ANOVA test to detect the difference between the curves. The significant differences at each point were calculated by the estimated marginal means (EMMeans), corrected by Tukey’s method. Lme4 v1.1.36, car v3.1.3, lmerTest v3.1.3 packages were used for these analyses. For statistical analysis for histograms, non-parametric Kruskal-Wallis tests were first used to assess overall differences among groups, followed by non-parametric, unpaired two-tailed Mann-Whitney tests for pairwise comparisons (*α* = 5%). These analysis were performed using GraphPad 6.01 Prism for Windows (GraphPad Software http://www.graphpad.com).

## 5 Data Availability

The datasets generated during the current study are available in the Zenodo repository, https://doi.org/10.5281/zenodo.16220127.

## 6 Acknowledgments

This study received grants from European Union’s MSCA grant MEDMOTILUS project (grant N°101024357), The National Agency of Research ANR PushPull (ANR-22-CE30), the PEPR microbiota JEMINI, INSERM IRP (Parasum), and Institut Carnot France Futur Elevage (“OrganoPig” project)”. Axel Ranson is recipient of a Ph.D. Fellowship from Ecole Normale Supérieure (ENS). The authors acknowldege Corinne Lencina (UMR GenPhySE, INRAE, Toulouse) for the quantification of IgA in mucus. EC thanks the Institut Universitaire de France (IUF) for support. Work in the Gorochov lab is supported by Institut National de la Santé et de la Recherche Médicale (INSERM), Sorbonne Université and the French Foundation for Medical Research (FRM) Paris, France (FRM EQU202203014622).

## 7 Author contributions

Conception & design: TLR, KC and EC. Resources: KS, RB, TLR, MVG, DS, GG, MB, EC. Experimental Data production KS, RB, TLR, MVG. Statistical analysis: KS, AR. Technical & material support: DS, GG. Supervision: KC and EC. Writing-original draft: KS, AR, KC and EC. All authors have read and approved the current version of the manuscript.

## Supplementary information

**Table S1:** Statistics for the main figure 1e.

| Contrast | Time | Estimate | SE | df | t.ratio | p.value | signif |
| --- | --- | --- | --- | --- | --- | --- | --- |
| cont - test | 0 | -3.4756 | 15.52958 | 16.75205 | -0.22381 | 0.825616 | ns |
| cont - test | 2 | -10.191 | 15.52958 | 16.75205 | -0.65623 | 0.520589 | ns |
| cont - test | 4 | -19.2096 | 15.52958 | 16.75205 | -1.23697 | 0.233159 | ns |
| cont - test | 6 | -19.2633 | 15.52958 | 16.75205 | -1.24042 | 0.231912 | ns |
| cont - test | 8 | -17.8121 | 15.52958 | 16.75205 | -1.14698 | 0.267508 | ns |
| cont - test | 10 | -25.2359 | 15.52958 | 16.75205 | -1.62502 | 0.122822 | ns |
| cont - test | 12 | -26.275 | 15.52958 | 16.75205 | -1.69194 | 0.109174 | ns |
| cont - test | 14 | -30.8762 | 15.52958 | 16.75205 | -1.98822 | 0.063382 | ns |
| cont - test | 16 | -30.4614 | 15.52958 | 16.75205 | -1.96151 | 0.066658 | ns |
| cont - test | 18 | -40.6413 | 15.52958 | 16.75205 | -2.61703 | 0.018186 | * |
| cont - test | 20 | -40.8009 | 15.52958 | 16.75205 | -2.6273 | 0.017805 | * |
| cont - test | 22 | -45.2921 | 15.52958 | 16.75205 | -2.9165 | 0.00973 | ** |
| cont - test | 24 | -47.2123 | 15.52958 | 16.75205 | -3.04015 | 0.007488 | ** |
| cont - test | 26 | -53.8156 | 15.52958 | 16.75205 | -3.46536 | 0.003012 | ** |
| cont - test | 28 | -50.1198 | 15.52958 | 16.75205 | -3.22738 | 0.005022 | ** |
| cont - test | 30 | -54.9198 | 15.52958 | 16.75205 | -3.53647 | 0.002584 | ** |
| cont - test | 32 | -52.3845 | 15.52958 | 16.75205 | -3.37321 | 0.003672 | ** |
| cont - test | 34 | -57.9909 | 15.52958 | 16.75205 | -3.73422 | 0.001687 | ** |
| cont - test | 36 | -60.4345 | 15.52958 | 16.75205 | -3.89158 | 0.001202 | ** |
| cont - test | 38 | -61.1668 | 15.52958 | 16.75205 | -3.93873 | 0.001086 | ** |
| cont - test | 40 | -61.2528 | 15.52958 | 16.75205 | -3.94426 | 0.001073 | ** |
| cont - test | 42 | -68.4598 | 15.52958 | 16.75205 | -4.40835 | 0.000397 | *** |
| cont - test | 44 | -71.5307 | 15.52958 | 16.75205 | -4.60609 | 0.000261 | *** |
| cont - test | 46 | -70.3044 | 15.52958 | 16.75205 | -4.52713 | 0.000308 | *** |
| cont - test | 48 | -71.787 | 15.52958 | 16.75205 | -4.6226 | 0.000252 | *** |
| cont - test | 50 | -70.9786 | 15.52958 | 16.75205 | -4.57054 | 0.000281 | *** |
| cont - test | 52 | -75.2617 | 15.52958 | 16.75205 | -4.84634 | 0.000158 | *** |
| cont - test | 54 | -75.4106 | 15.52958 | 16.75205 | -4.85593 | 0.000154 | *** |
| cont - test | 56 | -76.5601 | 15.52958 | 16.75205 | -4.92995 | 0.000132 | *** |
| cont - test | 58 | -75.9575 | 15.52958 | 16.75205 | -4.89115 | 0.000143 | *** |
| cont - test | 60 | -78.3004 | 15.52958 | 16.75205 | -5.04201 | 0.000105 | *** |
| cont - test | 62 | -76.7241 | 15.52958 | 16.75205 | -4.94051 | 0.000129 | *** |
| cont - test | 64 | -79.1773 | 15.52958 | 16.75205 | -5.09848 | 9.33E-05 | **** |
| cont - test | 66 | -77.9062 | 15.52958 | 16.75205 | -5.01664 | 0.000111 | *** |
| cont - test | 68 | -77.4841 | 15.52958 | 16.75205 | -4.98946 | 0.000117 | *** |
| cont - test | 70 | -79.2561 | 15.52958 | 16.75205 | -5.10356 | 9.23E-05 | **** |
| cont - test | 72 | -79.7562 | 15.52958 | 16.75205 | -5.13576 | 8.64E-05 | **** |
| cont - test | 74 | -80.0714 | 15.52958 | 16.75205 | -5.15606 | 8.29E-05 | **** |
| cont - test | 76 | -80.6268 | 15.52958 | 16.75205 | -5.19182 | 7.7E-05 | **** |
| cont - test | 78 | -80.7596 | 15.52958 | 16.75205 | -5.20037 | 7.57E-05 | **** |
| cont - test | 80 | -80.3783 | 15.52958 | 16.75205 | -5.17582 | 7.96E-05 | **** |
| cont - test | 82 | -79.9657 | 15.52958 | 16.75205 | -5.14925 | 8.4E-05 | **** |

Table S1: Statistics for the main figure 1e (Continuation)
| Contrast | Time | Estimate | SE | df | t.ratio | p.value | signif |
| --- | --- | --- | --- | --- | --- | --- | --- |
| cont - test | 84 | -81.0969 | 15.52958 | 16.75205 | -5.22209 | 7.24E-05 | **** |
| cont - test | 86 | -81.1656 | 15.52958 | 16.75205 | -5.22651 | 7.17E-05 | **** |
| cont - test | 88 | -81.1005 | 15.52958 | 16.75205 | -5.22232 | 7.23E-05 | **** |
| cont - test | 90 | -80.865 | 15.52958 | 16.75205 | -5.20716 | 7.46E-05 | **** |
| cont - test | 92 | -80.5113 | 15.52958 | 16.75205 | -5.18439 | 7.82E-05 | **** |
| cont - test | 94 | -81.5704 | 15.52958 | 16.75205 | -5.25258 | 6.8E-05 | **** |
| cont - test | 96 | -80.7937 | 15.52958 | 16.75205 | -5.20257 | 7.53E-05 | **** |
| cont - test | 98 | -80.1273 | 15.52958 | 16.75205 | -5.15966 | 8.23E-05 | **** |
| cont - test | 100 | -79.5621 | 15.52958 | 16.75205 | -5.12326 | 8.87E-05 | **** |
| cont - test | 102 | -79.811 | 15.52958 | 16.75205 | -5.13929 | 8.58E-05 | **** |
| cont - test | 104 | -80.6662 | 15.52958 | 16.75205 | -5.19436 | 7.66E-05 | **** |
| cont - test | 106 | -79.9791 | 15.52958 | 16.75205 | -5.15012 | 8.39E-05 | **** |
| cont - test | 108 | -79.8019 | 15.52958 | 16.75205 | -5.13871 | 8.59E-05 | **** |
| cont - test | 110 | -79.1715 | 15.52958 | 16.75205 | -5.09811 | 9.34E-05 | **** |
| cont - test | 112 | -78.5389 | 15.52958 | 16.75205 | -5.05738 | 0.000102 | *** |
| cont - test | 114 | -79.1217 | 15.52958 | 16.75205 | -5.0949 | 9.4E-05 | **** |
| cont - test | 116 | -77.7565 | 15.52958 | 16.75205 | -5.00699 | 0.000113 | *** |
| cont - test | 118 | -77.5875 | 15.52958 | 16.75205 | -4.99611 | 0.000115 | *** |
| cont - test | 120 | -78.14 | 15.52958 | 16.75205 | -5.03169 | 0.000107 | *** |
| cont - test | 122 | -77.4989 | 15.52958 | 16.75205 | -4.9904 | 0.000117 | *** |
| cont - test | 124 | -76.8447 | 15.52958 | 16.75205 | -4.94828 | 0.000127 | *** |
| cont - test | 126 | -76.4769 | 15.52958 | 16.75205 | -4.9246 | 0.000134 | *** |
| cont - test | 128 | -76.0418 | 15.52958 | 16.75205 | -4.89658 | 0.000142 | *** |
| cont - test | 130 | -75.483 | 15.52958 | 16.75205 | -4.8606 | 0.000153 | *** |
| cont - test | 132 | -75.9101 | 15.52958 | 16.75205 | -4.8881 | 0.000144 | *** |
| cont - test | 134 | -75.4679 | 15.52958 | 16.75205 | -4.85962 | 0.000153 | *** |
| cont - test | 136 | -75.4415 | 15.52958 | 16.75205 | -4.85792 | 0.000154 | *** |
| cont - test | 138 | -75.1085 | 15.52958 | 16.75205 | -4.83648 | 0.000161 | *** |
| cont - test | 140 | -74.5164 | 15.52958 | 16.75205 | -4.79835 | 0.000174 | *** |
| cont - test | 142 | -72.9611 | 15.52958 | 16.75205 | -4.6982 | 0.000215 | *** |
| cont - test | 144 | -72.651 | 15.52958 | 16.75205 | -4.67824 | 0.000224 | *** |
| cont - test | 146 | -72.7229 | 15.52958 | 16.75205 | -4.68286 | 0.000222 | *** |
| cont - test | 148 | -72.9191 | 15.52958 | 16.75205 | -4.6955 | 0.000216 | *** |

The significant differences at each time were calculated by the estimated marginal means (EMMeans), corrected by Tukey's method. ns (not significant): $p > 0.05$ , \*: $p \leq 0.05$ , \*\*: $p \leq 0.01$ , \*\*\*: $p \leq 0.001$ , \*\*\*\*: $p \leq 0.0001$ .

**Table S2:** Statistics for the main figure 2b.

| contrast | r/R | estimate | SE | df | t.ratio | p.value | signif |
| --- | --- | --- | --- | --- | --- | --- | --- |
| cont - test | 0.1 | -1E-05 | 0.000173 | 346.4056 | -0.05961 | 0.952504 | ns |
| cont - test | 0.2 | 1.6E-05 | 0.000129 | 327.418 | 0.123916 | 0.901457 | ns |
| cont - test | 0.3 | -9.6E-06 | 0.000119 | 315.7509 | -0.08046 | 0.935923 | ns |
| cont - test | 0.4 | -2E-05 | 0.000115 | 308.9307 | -0.17401 | 0.861968 | ns |
| cont - test | 0.5 | -4.1E-05 | 0.000115 | 308.9307 | -0.35428 | 0.723368 | ns |
| cont - test | 0.6 | -0.00012 | 0.000115 | 308.9307 | -1.01267 | 0.312011 | ns |
| cont - test | 0.7 | -0.0003 | 0.000115 | 308.9307 | -2.63252 | 0.008901 | ** |
| cont - test | 0.8 | -0.00066 | 0.000115 | 308.9307 | -5.76685 | 1.96E-08 | **** |
| cont - test | 0.9 | -0.00126 | 0.000115 | 308.9307 | -10.9764 | 6.9E-24 | **** |
| cont - test | 1 | -0.00065 | 0.000115 | 308.9307 | -5.64801 | 3.68E-08 | **** |
| cont - test | 1.1 | 0.00085 | 0.000115 | 308.9307 | 7.41767 | 1.16E-12 | **** |
| cont - test | 1.2 | 3.37E-05 | 0.000115 | 308.9307 | 0.29433 | 0.768703 | ns |
| cont - test | 1.3 | 2.85E-06 | 0.000115 | 308.9307 | 0.024874 | 0.980171 | ns |
| cont - test | 1.4 | 3.02E-05 | 0.000115 | 308.9307 | 0.263859 | 0.792064 | ns |
| cont - test | 1.5 | 7.32E-05 | 0.000115 | 308.9307 | 0.638761 | 0.523452 | ns |
| cont - test | 1.6 | 7.46E-05 | 0.000115 | 308.9307 | 0.65067 | 0.515743 | ns |
| cont - test | 1.7 | 9.04E-05 | 0.000115 | 308.9307 | 0.788366 | 0.431087 | ns |
| cont - test | 1.8 | 6.52E-05 | 0.000115 | 308.9307 | 0.569092 | 0.569707 | ns |
| cont - test | 1.9 | 5.94E-05 | 0.000115 | 308.9307 | 0.518402 | 0.604549 | ns |
| cont - test | 2 | 6.24E-05 | 0.000115 | 308.9307 | 0.54485 | 0.58625 | ns |
| cont - test | 2.1 | 5.01E-05 | 0.000115 | 308.9307 | 0.437173 | 0.662291 | ns |
| cont - test | 2.2 | 4.66E-05 | 0.000115 | 308.9307 | 0.40631 | 0.684796 | ns |
| cont - test | 2.3 | 2.55E-05 | 0.000115 | 308.9307 | 0.222885 | 0.823772 | ns |
| cont - test | 2.4 | 2.09E-05 | 0.000115 | 308.9307 | 0.182372 | 0.855411 | ns |
| cont - test | 2.5 | 1.96E-05 | 0.000115 | 308.9307 | 0.171404 | 0.864019 | ns |
| cont - test | 2.6 | 7.09E-06 | 0.000115 | 308.9307 | 0.061872 | 0.950705 | ns |
| cont - test | 2.7 | -1.6E-05 | 0.000115 | 308.9307 | -0.13648 | 0.891529 | ns |
| cont - test | 2.8 | -3E-05 | 0.000115 | 308.9307 | -0.2584 | 0.796271 | ns |
| cont - test | 2.9 | -2.4E-05 | 0.000115 | 308.9307 | -0.20742 | 0.835818 | ns |
| cont - test | 3 | -3E-05 | 0.000119 | 315.8225 | -0.24787 | 0.8044 | ns |

Significant differences at each r/R were calculated by the estimated marginal means (EMMeans), and corrected by Tukey's method. ns (not significant): $p > 0.05$ , \*\*: $p \leq 0.01$ , \*\*\*\*: $p \leq 0.0001$ .

**Fig. S1.**
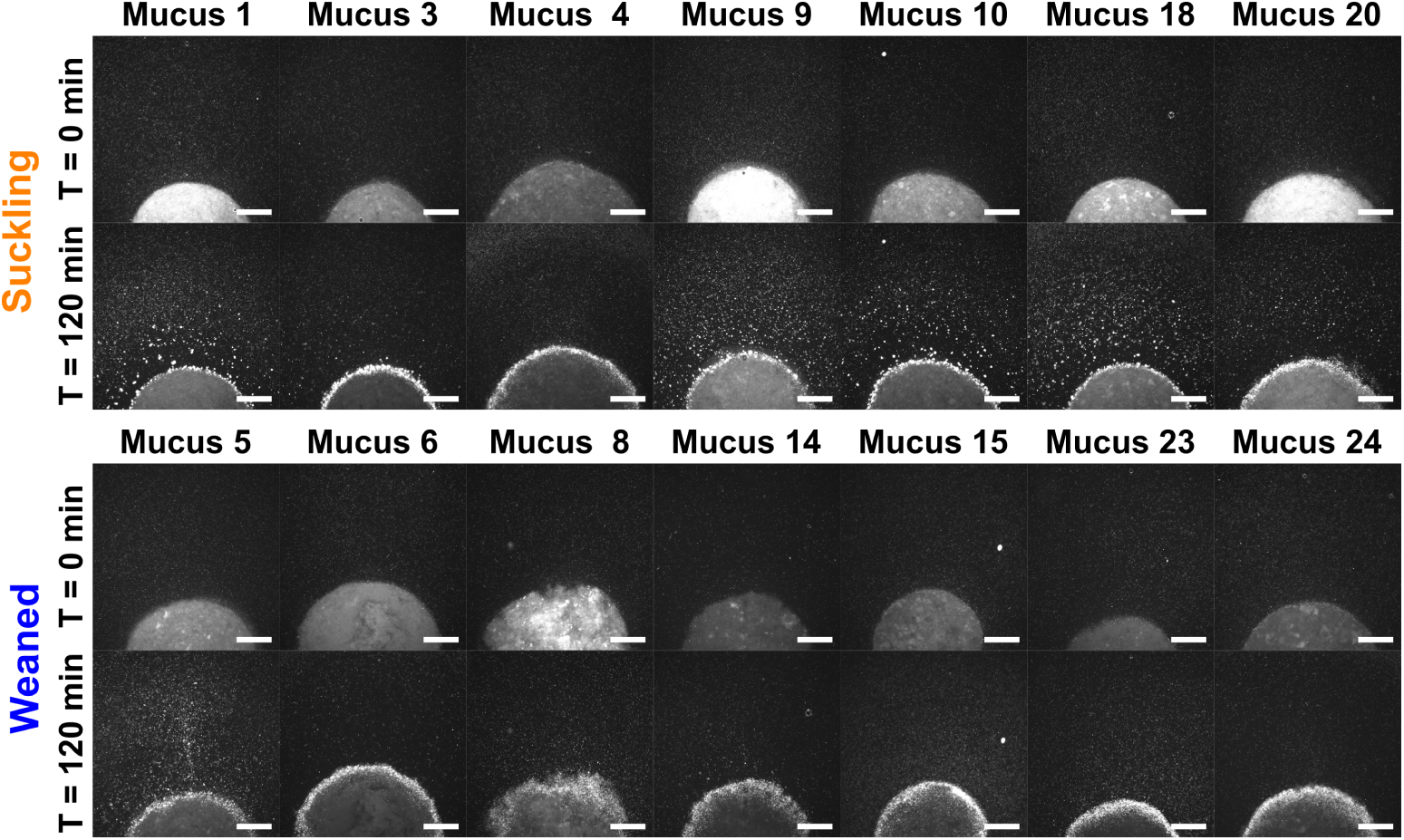
Time-lapse experiments of droplets from the mucus samples used in this work. Images at time-points *T* = 0 min and *T* = 120 min from time-lapse experiments of *E. coli* (bright dots) in a mucus droplet from a suckling or a weaned piglet. The figure shows representative images of droplets from all the mucus samples used in this work to calculate the penetration length and the aggregation of bacteria (*n* = 7 per group). Scale bars 500 *µ*m.

**Fig. S2.**
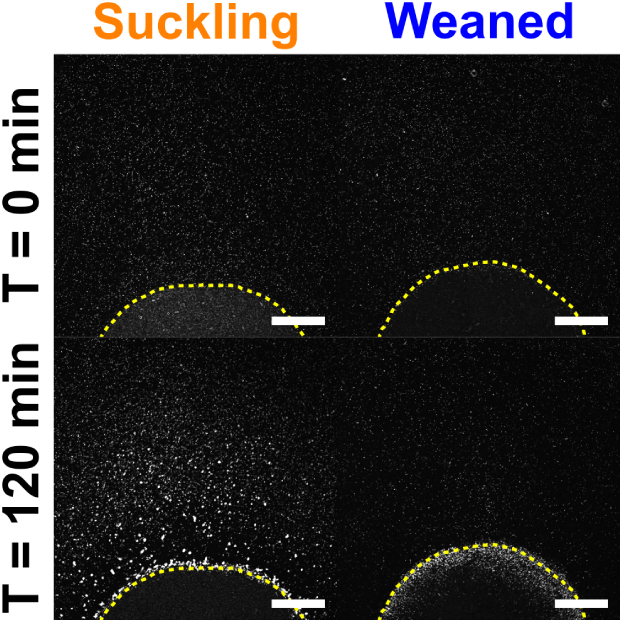
Delimitation of the edges of mucus droplets. First (*T* = 0 min) and last (*T* = 120 min) images of a time-lapse experiment showing the interface between a mucus droplet from suckling and weaned piglets and a bacterial suspension (white dots). The yellow dashed lines represent the edge of the mucus droplets, which was manually defined at *T* = 0 min, and they divide the images into mucus exterior and mucus interior. Scale bars 500 *µ*m.

**Fig. S3.**
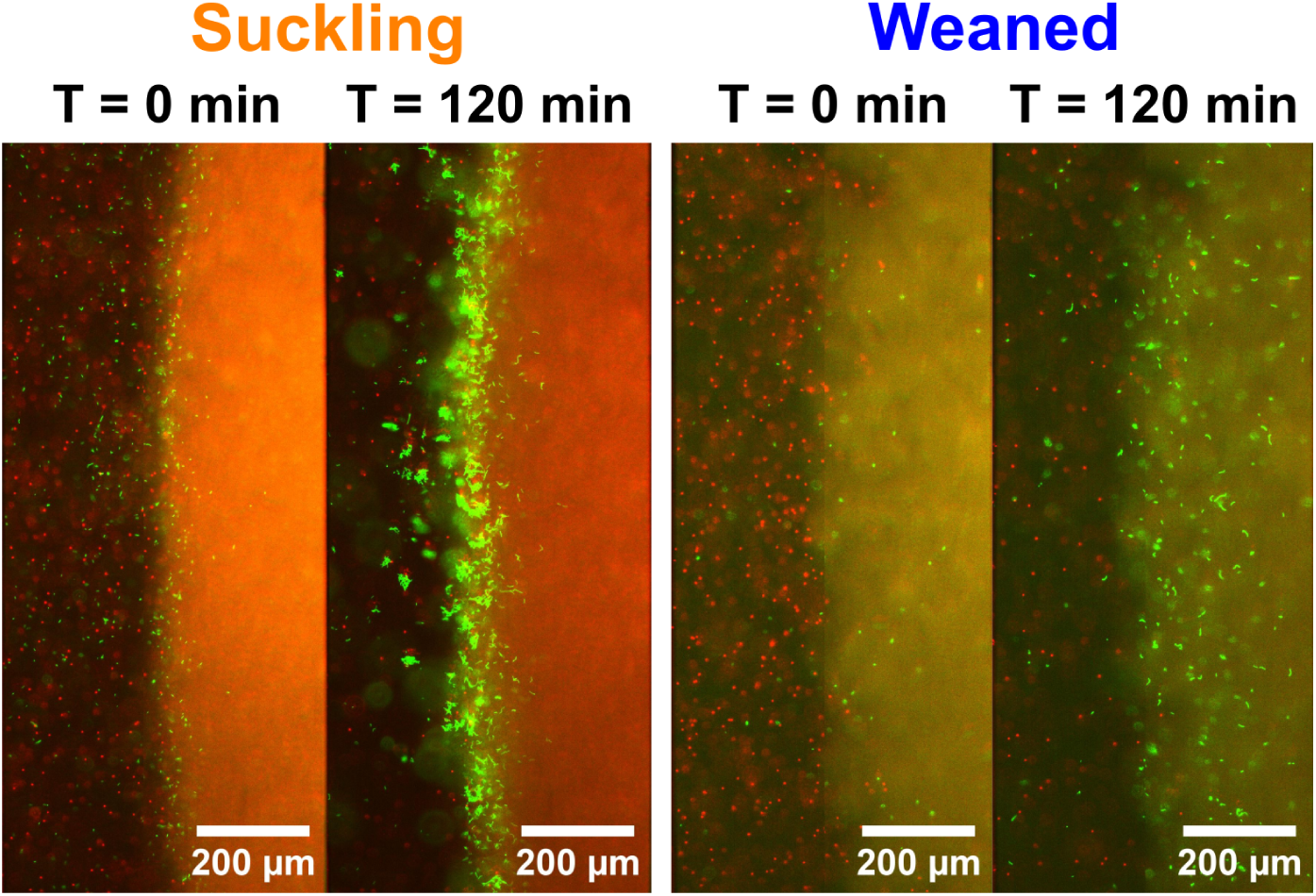
Motility is needed to penetrate the mucus layer. First (*T* = 0 min) and last (*T* = 120 min) images of a time-lapse experiment showing the interface between mucus from suckling and weaned piglets and a bacterial suspension. Passive tracers (1 *µ*m fluorescent beads, red dots) were included in the bacterial suspension to show that they cannot invade the mucus layer whereas the green fluorescent body of motile bacteria can be identified in the mucus layer.

**Fig. S4.**
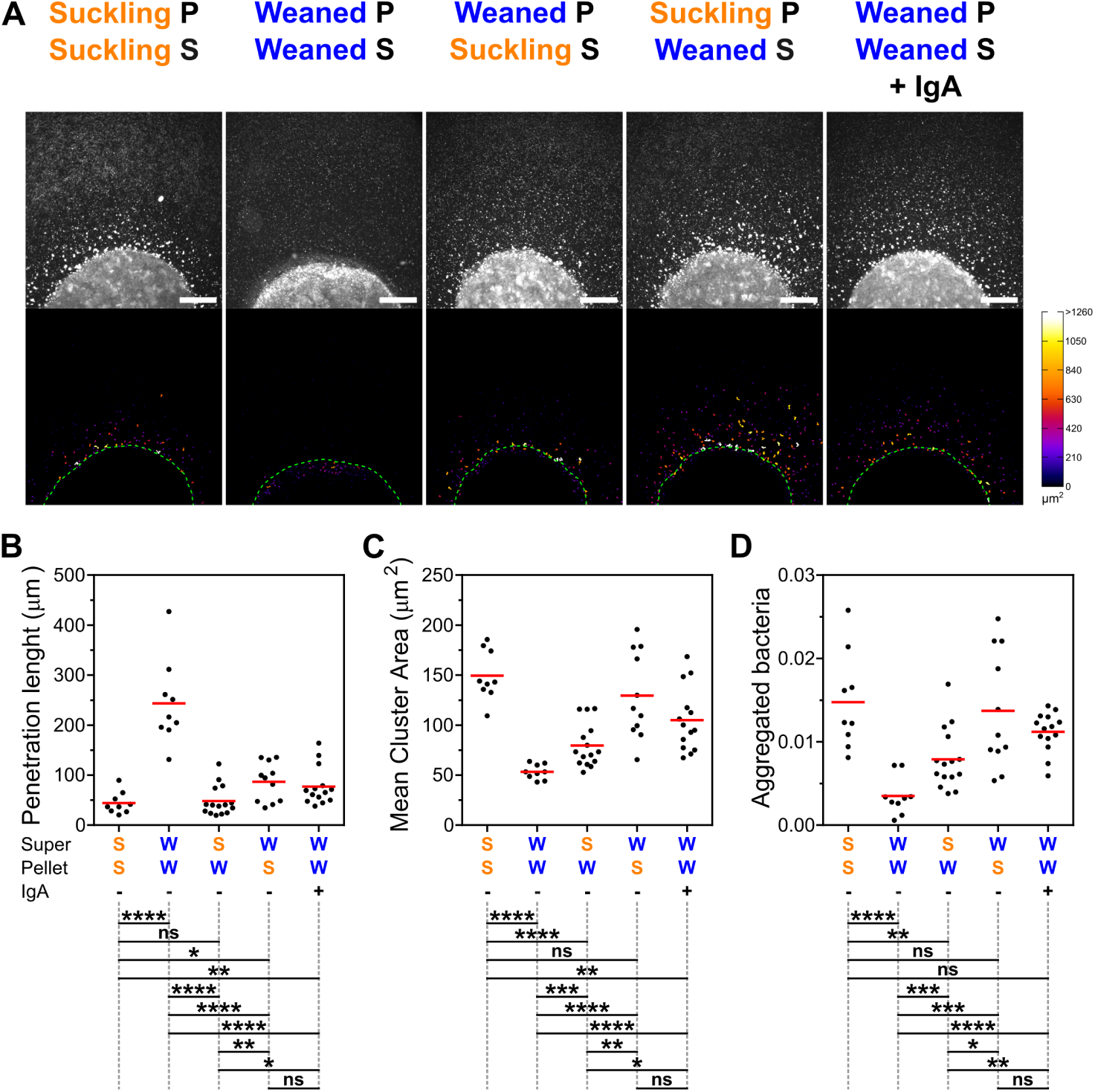
Complete set of supernatant/pellet combinations. (A) Representative images of mucus droplets obtained by mixing supernatant (S) and pellet (P) from suckling and weaned mucus. Droplets were introduced into the microfluidic system which was then filled with a bacterial suspension, and images were acquired at *T* = 120 min. Bottom panels show binarized views of the top panels, with bacterial aggregates color-coded according to their area. Green dashed lines delimitate mucus boundaries. Scale bars 500 *µ*m. (B-D) Quantification of bacterial penetration length (B), mean area of aggregated bacteria (C), and total fraction of aggregated bacteria (D) of mucus droplets composed of different supernatant/pellet combinations from suckling (S) and weaned (W) mucus. The penetration length was calculated using the data from the interior region, while the mean area and total fraction of aggregated bacteria were calculated for the external region. Each dot represents and individual experiment. Red horizontal lines denote group means. The table below each graph show the statistical differences based on non-parametric, unpaired two-tailed Mann-Whitney test (*α* = 5%). ns (not significant): p *>* 0.05, ∗: p ≤ 0.05, ∗∗: p ≤ 0.01, ∗ ∗ ∗: p ≤ 0.001, ∗ ∗ ∗∗: p ≤ 0.0001.

**Fig. S5.**
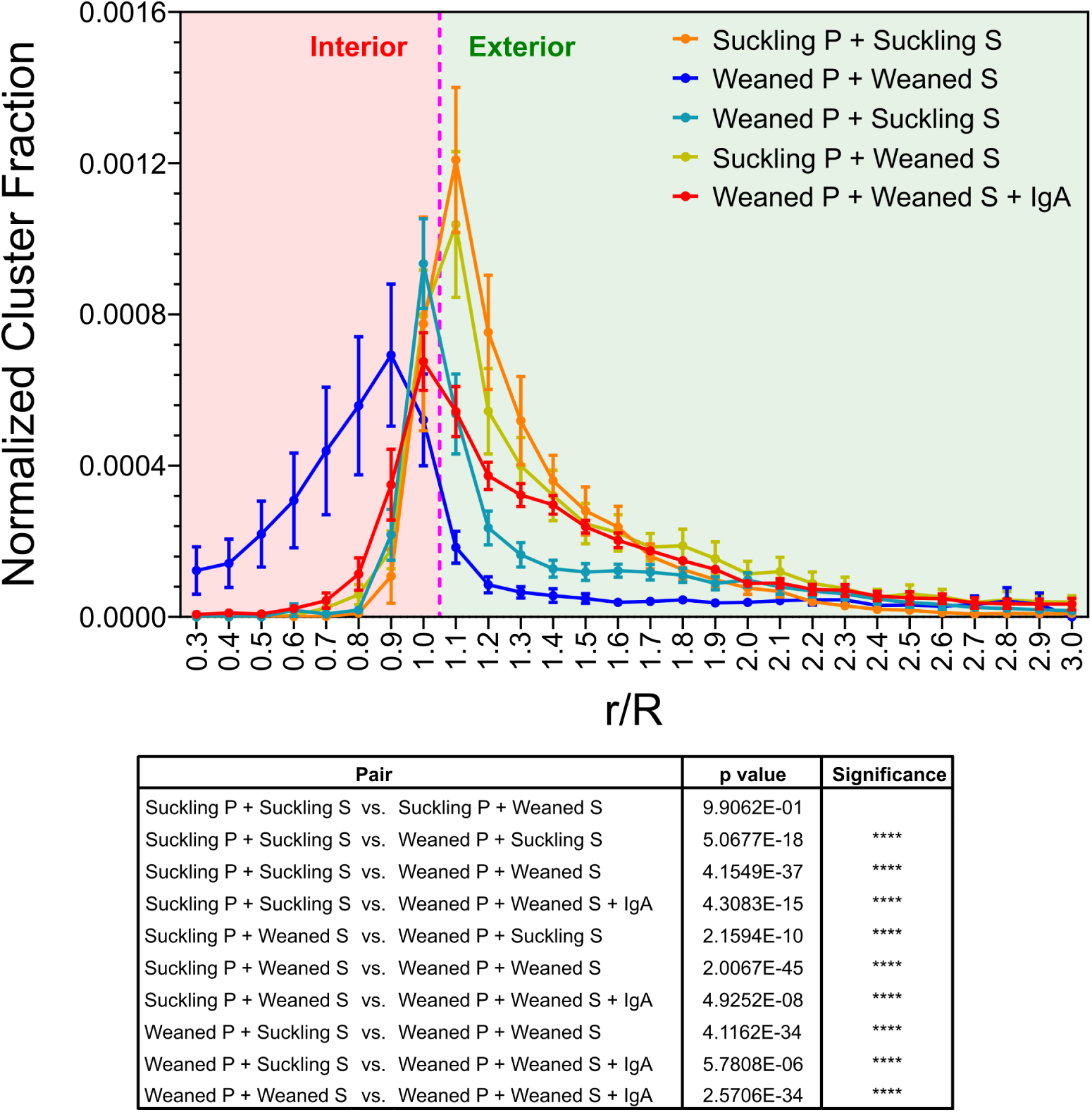
Fraction of aggregated bacteria per subregion in supernatant/pellet combinations. Quantification of the normalized fraction of aggregated bacteria at *T* = 120 min across defined subregions of mucus droplets composed of different supernatant (S) and pellet (P) combinations. Data represent the mean ± standard error of the mean (SEM). Number of replicates: Suckling P + Suckling S: 9, Weaned P + Weaned S: 9, Weaned P + Suckling S: 15, Suckling P + Weaned S: 11, Weaned P + Weaned S + IgA: 14. Magenta dashed line: Mucus boundary. The results of type III ANOVAs performed on linear mixed models derived from these data, 2 groups at 2, are shown below the graph.

